# Extrahepatic effects of thyroid hormone and resmetirom override their beneficial hepatic effects in alcohol-associated liver disease in mice

**DOI:** 10.1101/2025.10.10.681680

**Authors:** Christoph Hoppe, G. Sebastian Hönes, Adrian D. Prinz, Devon Siemes, Christina Wenzek, Susanne C. Grund, Ann-Kathrin Schörding, Yara M. Machlah, Eveline Bruinstroop, David Laehnemann, Manuel Philipp, Karine Gauthier, Frédéric Flamant, Daniel R. Engel, Hideo A. Baba, Johannes Köster, Dagmar Führer, Denise Zwanziger, Christian M. Lange, Lars C. Moeller

**Affiliations:** Department of Endocrinology, Diabetes and Metabolism and Division of Laboratory Research, University Hospital Essen, Essen, Germany; Bioinformatics and Computational Oncology, Institute for Artificial Intelligence in Medicine, University Hospital Essen, University of Duisburg-Essen, Essen, Germany; Institute of Experimental Immunology and Imaging, University Hospital Essen, Essen, Germany; Department of Endocrinology and Metabolism, Amsterdam UMC, University of Amsterdam, Amsterdam, Netherlands; Institut de Génomique Fonctionnelle de Lyon, École normale supérieure de Lyon, Lyon, France; Institute of Pathology, University Hospital Essen, Essen, Germany; Department of Internal Medicine II, Ludwig-Maximilian-University Hospital Munich, Munich, Germany

**Keywords:** thyroid hormone, liver, ALD, resmetirom, liver-WAT-axis, thyroid hormone receptor β signaling

## Abstract

**Background:** Alcohol-associated liver disease (ALD) is a common type of liver disease worldwide. Excessive consumption of ethanol (EtOH) causes fat accumulation leading to hepatic steatosis. Hepatic thyroid hormone (TH) action or liver-specific thyromimetics, e.g. resmetirom, can reduce hepatic triglycerides. We therefore hypothesized that TH treatment could ameliorate ALD.

**Methods:** To induce ALD, mice were treated with either EtOH or liquid control diet for 10 days followed by a single EtOH or maltose control gavage on day 11. The liquid diets were supplemented with solvent, T3 or Resmetirom. We studied WT and hepatocyte-specific TRβ KO mice (hepTRβKO). Effects were measured by clinical chemistry, liver staining, hepatic triglyceride content, and RNA-sequencing.

**Results:** Surprisingly, resmetirom had no beneficial effect and T3 treatment even aggravated EtOH-induced steatosis (increased liver weight and hepatic triglycerides). The liver phenotype was worsened in hepTRβKO mice, which still suggested beneficial effects of hepatic TRβ signaling in WT mice. These seemingly paradoxical results could be explained by extrahepatic effects in WAT: WAT weight and adipocyte size were reduced by EtOH, T3 and resmetirom. These data indicate lipolysis and subsequent fatty acid accumulation in the liver, explaining the more severe ALD phenotype with T3 and attenuated effect of resmetirom. As WAT loss was reduced in hepTRβKO mice, the hepatic TRβ mediated the extrahepatic effects of T3 and resmetirom on WAT.

**Conclusion:** We conclude that extrahepatic TH effects in WAT were detrimental in ALD and counteracted beneficial local hepatic TH/TRβ action. As WAT loss appeared to originate from the hepatic TRβ, this also applied to resmetirom.

## Introduction

Thyroid hormone (TH) and TH receptor (TR) β action promote lipid catabolism in the liver and reduce hepatic lipid content. This is especially apparent in situations of impaired TH/TRβ signaling. In mice with a TRβ mutation that prevents TH binding, hepatic lipid content was increased.^1^ This was attributed to increased expression of lipogenic genes. Similarly, we found increased liver triglycerides (TG) in TRβ knockout (KO) mice compared to their wildtype (WT) littermates.^2^ These results from mice could be confirmed in humans. Patients with resistance to TH due to a mutation in the *THRB* gene (RTHβ) had increased hepatic fat content compared to their unaffected relatives.^3^ While the hepatic fat content in unaffected relatives was strongly correlated to age and BMI, this correlation was absent in the RTHβ patients, indicating that TH/TRβ signaling in liver is predominant over the effects of age and obesity.

While TRβ signaling in hepatocytes acts simultaneously to enhance both lipogenesis and fatty acid oxidation,^4^ the latter mechanism is most relevant and determines the net TH effect of reduced fat content.^5^ For these reasons, therapeutic application of TH action to reduce hepatic steatosis is desirable. In mice, TH treatment prevented hepatic steatosis on a nonalcoholic steatohepatitis (NASH) diet,^6^ demonstrating its efficacy. To avoid detrimental TH effects outside the liver, e.g. tachycardia, atrial fibrillation or osteoporosis, liver-directed and TRβ-specific TH analogs like resmetirom have been developed. We found that resmetirom uses a hepatocyte-specific transporter (OATP1B1) for entry into cells, which may explain its liver specificity, and that resmetirom indeed predominantly acts on TRβ.^7^

Resmetirom has been studied in clinical phase III trials for treatment of NASH, now called Metabolic dysfunction-associated steatohepatitis, (MASH). It met the goals set by the FDA, briefly improvement of fibrosis without worsening of NASH score or improvement of the NASH score without worsening of fibrosis.^8^ Based on these data, resmetirom was conditionally approved by the FDA in March 2024. Subsequently, resmetirom was recommended for treatment of adults with non-cirrhotic MASH and significant liver fibrosis (stage ≥2).^9^

Alcohol-associated liver disease (ALD) is a major cause of chronic liver disease worldwide. ALD ranges from simple steatosis to life-threatening steatohepatitis, finally cirrhosis, and superimposed hepatocellular carcinoma. The clinically relevant question was therefore: Is TH action and especially resmetirom also beneficial inALD? To answer this question, we treated mice with EtOH following the NIAAA protocol to induce ALD in male mice^10^ and studied the effects of concomitant T3 or resmetirom treatment.

### Materials and Methods

#### Animals and Treatment

All animal studies were approved by the local authorities (Landesamt für Natur, Umwelt und Verbraucherschutz Nordrhein-Westfalen). Wildtype (WT, TRβ^+/fl^, Alb-Cre^0/tg^), hepatocyte-specific TRβ knockout (hepTRβKO, TRβ^-/fl^, Alb-Cre^0/tg^) and hepatocyte-specific TRβ mutant (hepTRβGS, TRβ^GS/fl^, Alb-Cre^0/tg^) mice were generated by crossing Thrb-floxed mice with heterozygous Thrb knockout or mutant mice expressing Cre recombinase under the control of the mouse albumin promoter. Only male mice were studied. Mice were housed at 22 ± 2 °C under a 12-hour light/12-hour dark cycle. All cages contained nesting material, a balcony and wood. Mice (8-10 weeks old) were subjected to chronic and binge feeding of EtOH as established in the NIAAA (National Institute of Alcohol Abuse and Alcoholism) model by Bertola et al.^10^ Mice received a liquid diet [Lieber-DeCarli (LDC) Shake and Pour control liquid diet, Bio-Serv, F1259SP] for acclimatization for 5 days. Mice were then assigned to different treatment groups: LDC, LDC+T3, LDC+Resmetirom as control and EtOH, EtOH+T3 and EtOH+Resmetirom, receiving Lieber-DeCarli ethanol (EtOH) liquid diet (5 % ethanol, Bio-Serv, F1258SP) (Figure 1A). Liquid diets were given for the next 11 days with daily food exchange in liquid diet feeding tubes (Bio-Serv, 9019, 50 mL). EtOH liquid diet was available *ad libitum*, while the control diet was pair-fed. Weight and appearance of mice were monitored daily during the experiment. On day 11, mice received an oral gavage with either ethanol (5 g ethanol/kg BW) or maltose (9 g maltose dextrin/kg BW, Bio-Serv, 10DE, 3585) according to their liquid diet. After 9 hours, mice were sacrificed, and blood and tissue samples were collected.

**Figure 1.**
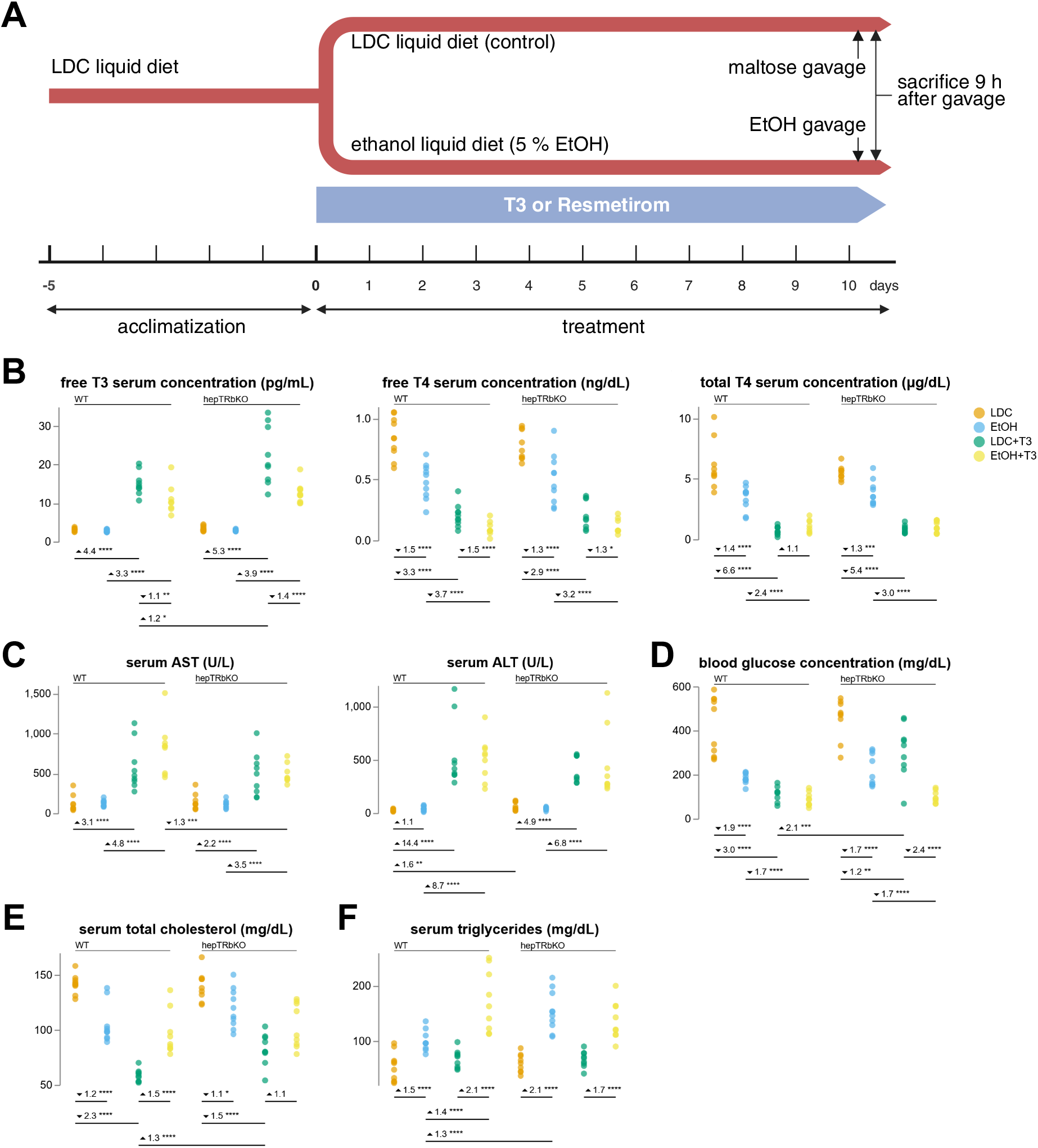
Effects of EtOH and T3 treatment on serum parameter in ALD. (A) experimental overview of liquid control diet and EtOH diet feeding, (B) free T3, free T4 and total T4 serum concentrations, (C) serum levels of aspartate transaminase (AST) and alanine transaminase (ALT), (D) blood glucose concentration, (E) total cholesterol serum concentrations and (F) serum triglyceride levels. CFC, BM:*P<0.05, **P<0.01, ***P<0.001, ****P<0.0001. n=7-10.

For TH treatment, liquid diets were supplemented with T3 (100 µg/kg BW, Sigma-Aldrich, T6397) or resmetirom (10 mg/kg BW, MedChemExpress, HY-12216) for the entire 11 days.

### Serum Measurements and Thyroid Function Tests

Blood was collected by cardiac puncture. Blood glucose levels were measured immediately after collecting with Accu-Chek Guide glucometer (Roche) and test stripes. Blood samples were centrifuged to obtain serum samples. Serum aspartate and alanine transaminase levels (AST/ALT), serum total cholesterol (TC) and serum triglycerides (TG) concentrations were measured using Spotchem™ EZ SP-4430(Arkray). Serum free triiodothyronine (FT3), free thyroxine (FT4) and total thyroxine (TT4) concentrations were determined by ELISA according to manufacturer’s instructions (EIA-2385, EIA-2386, EIA-1781, DRG Instruments, Germany).

### Hepatic Triglycerides

Hepatic triglycerides were extracted from liver (50-100 mg) with chloroform:methanol (2:1) for 2 hours at room temperature in an orbital shaker. Supernatants were washed with 0.9 % NaCl (B.Braun). The organic phase was transferred after centrifugation and dried under vacuum. The extracted triglycerides were dissolved in 2 % Triton X-100 (Sigma, T8787) and measured with a triglyceride colorimetric assay kit (Cayman) using a standard curve.

### RNA Isolation and Gene Expression Analysis

Total RNA was isolated from 50 mg of homogenized liver following the manufacturer’s instructions (RNeasy Mini Kit, QIAGEN). Concentration and quality of RNA was measured with Agilent Bioanalyzer Nano (Agilent, Santa Clara, CA USA). RNA (600 ng) was reverse transcribed into cDNA with SuperScript III (Invitrogen) and random hexamers. Quantitative PCR (qPCR) was performed on Lightcycler LC480 (Roche). Transcripts were quantified by PerfeCTa SYBR Green SuperMix Rox (QuantaBio). Relative mRNA levels were normalized to the expression of *Ppia* mRNA and calculated with efficiency correction.

For RNA-Sequencing, libraries were prepared with Lexogens QuantSeq 3’ mRNA-Seq Library Prep Kit FWD, quality controlled with Qubit (Invitrogen, Waltham, MA, USA) and library quant qPCR, and sequenced on a NextSeq500 75 cycles (Illumina, San Diego, CA, USA). Data were analyzed with the rna-seq-kallisto-sleuth workflow (https://github.com/snakemake-workflows/rna-seq-kallisto-sleuth). Specifically, sequences were trimmed with Cutadapt.^11^ Transcripts were quantified while accounting for 3’ mRNA-Seq by mitigating the leaking of non-Matched Annotation of NCBI and EMBL (MANE) transcript expression into the MANE transcript expression estimates. This was achieved by mapping reads to the whole transcriptome using BWA-MEM^12^ keeping only reads that mapped closest to the 3’ end of the MANE transcript of each gene. Then, Kallisto^13^ was used to quantify the MANE transcripts with the filtered reads. Integrated normalization and transcript differential expression analysis was conducted with Sleuth,^14^ using the likelihood ratio test method while correcting for experimental cohort related batch effects. Sleuth effect size estimates (beta-scores) were calibrated to represent log2-fold changes by transforming input counts x with log2(x + 0.5). False discovery rate was controlled with the Benjamini-Hochberg method.^15^ Genes were sorted in a threshold-free approach utilizing the pi-value,^16^ which we reinterpreted to combine FDR with absolute values of sleuth beta-scores of the respective comparison variable, such that those genes with the strongest combination of significance and effect size are listed at the top. Gene ontology enrichment was performed on expressions of MANE transcripts (according to Ensembl Mus musculus release 111) using goatools,^17^ while defining transcripts with FDR<=0.05 as foreground. All results were visualized using Datavzrd (https://datavzrd.github.io) and Vega-Lite.^18^ Versions and parameter details as well as all workflow steps can be found in the Snakemake report (Supplement).

### H&EStaining and Adipocyte Cell Size

Tissues were fixed in 4 % paraformaldehyde for 24-48 hours and dehydrated in rising gradients of ethanol and xylol before paraffin embedding. Tissue sections (5 µm) were deparaffinized with Neo-Clear (Merck) and hydrated in ethanol (100 %, 96 % and 70 %). Sections were stained with hematoxylin (Mayer’s hemalaun) for 5 minutes, rinsed for 10 minutes with warm tap water and stained with eosin (Sigma Aldrich) for 45 seconds before repeated dehydration. Stained tissue sections were covered in Neo-Mount (Merck) and images were taken with upright epifluorescence microscope (Olympus BX51). Adipocyte cell size was determined with ImageJ and the plugin Adipocyte Tools.^19^

### Oil Red O Staining

Fresh liver samples were embedded in Tissue-Tek O.C.T. (Sakura Finetek) and frozen in the gas phase of liquid nitrogen. Three micrometer sections were cut and stained in Oil Red O solution for 10 minutes, rinsed in isopropanol (60%) and washed in distilled water. The sections were counterstained with hematoxylin for 1 minute, washed with running tap water and mounted with aqueous mounting medium. Images of the stained tissues were taken with Aperio ScanScope AT2 (Leica) and analyzed with Aperio Image Scope. The Oil Red O positive area was calculated within the whole tissue.

### Statistical Analysis of non RNA-seq measurements

To analyze the effect size between each pair of conditions in an uncertainty aware way, we conducted the following analysis. First, we empirically calculated the posterior distribution of the log2 fold change using bootstrapping. Specifically, for each pair of conditions under consideration, we resampled (while retaining the sample size) the measured values per condition 1000 times with replacement. For each such resampling, we calculated the log2 fold change. By binning and counting the resulting log2 fold changes across all 1000 repetitions, the empirical posterior distribution was obtained. From these, we obtained a conservative estimate of the fold change for reporting, by taking the log2 fold change closest to 0 (i.e. no change) such that 95% of the values in the empirical posterior distribution are more extreme. In other words, we anticipate the true effect size to be more extreme than the reported fold change with a probability of 95%. In the manuscript text and any plots, we present non log-space conservative fold changes (CFC), visualized with an up-pointing triangle for up-regulation, and a down-pointing triangle for down-regulation, complemented by the reciprocal value of the fold change (i.e. y/x instead of x/y, denoting an y/x-fold downregulation). In addition, we calculated the statistical significance via the non-parametric Brunner-Munzel (BM) test,^20^ obtaining a p-value for the null hypothesis that, for two randomly selected values x and y from the two compared conditions, the probability of x>y is equal to y>x. The p-values were adjusted for multiple testing using the Bonferroni-Holm method.^21^ The data plots show both the conservative fold change estimate, as well as the adjusted p-values (in star-notation, see respective figure captions) for selected comparisons of interest. In addition, we provide the conservative fold change of all pairwise condition comparisons per experiment and the associated empirical posterior distributions in the Snakemake report (Supplement). The entire analysis has been implemented as a generic reproducible Snakemake workflow (https://github.com/snakemake-workflows/effect-size-estimation). Additional plots were analyzed and visualized with Prism 10 (Graphpad Software).

## Results

### TH treatment and EtOH both severely aggravate metabolic serum parameters

To investigate the influence of TH signaling on ethanol-induced hepatic steatosis, we induced a mild chronic ALD in male WT and hepTRβKO mice by ethanol (EtOH) feeding based on the NIAAA mouse model (Figure 1A). T3 was administered for treatment in both the LDC control liquid diet and the EtOH liquid diet for 11 days. Serum TH concentrations (Figure 1B) were measured to confirm hyperthyroidism after T3 treatment. Because of exogenous T3 application, FT3 serum concentrations were increased, while FT4 and TT4 serum levels were decreased due to the negative feedback of the hypothalamic-pituitary-thyroid axis after T3 treatment. Noteworthy, EtOH treatment did not affect FT3 concentrations, but FT4 as well as TT4 serum concentrations were decreased (Figure 1B).

To assess the effect of TH treatment on ALD, we measured metabolic serum parameters in WT and hepTRβKO mice. Interestingly, both LDC+T3 and EtOH+T3 treatment increased serum AST and ALT concentrations to the same extent in WT as in hepTRβKO mice (Figure 1C). Total cholesterol (TC) concentrations were reduced by T3 in both LDC control groups, but more in WT mice. EtOH severely diminished this effect (Figure 1E). Triglyceride (TG) serum concentrations were increased by EtOH feeding with a stronger effect in hepTRβKO mice (Figure F). Surprisingly, in WT mice T3 treatment in combination with EtOH even further elevated TG concentration compared to the EtOH-only group (Figure 1F) while this was not the case for hepTRβKO mice. Contrary to our hypothesis, increased transaminase serum concentrations and elevated serum TG suggested that T3 treatment was rather unfavorable in ALD.

### TH treatment and absence of hepatic TRβ signaling aggravate EtOH-induced hepatic steatosis

Given these surprising results of unfavorable T3 effects on serum parameters in ALD, we determined whether local TH action in hepatocytes per se induces adverse or beneficial effects. To determine the extent of alcohol-associated hepatic steatosis, we measured liver weight and hepatic fat content. In WT mice, the liver/body weight ratio increased with EtOH and this effect was further pronounced with EtOH+T3 treatment (Figure 2A). Noteworthy, loss of hepatic TRβ in hepTRβKO mice aggravated this effect and, moreover, the liver/body weight ratio was also increased by T3 in the LDC group of hepTRβKO mice compared to WT. Similarly, hepatic TG content in hepTRβKO mice was strongly increased by T3 with LDC while there was no effect in WT mice treated with LDC and T3. However, the combination of T3 with EtOH led to two- to three-fold increased hepatic triglyceride content in both genotypes (Figure 2B). Importantly, Oil Red O staining confirmed that hepatic TG content was markedly increased by T3 treatment in hepTRβKO mice, independent from the diet (Figure 2C and D). These results demonstrate that T3 treatment in the absence of hepatic TRβ signaling is more detrimental, which, conversely, implies that local hepatic T3/TRβ action is beneficial in the liver and important for lipid metabolism.

**Figure 2.**
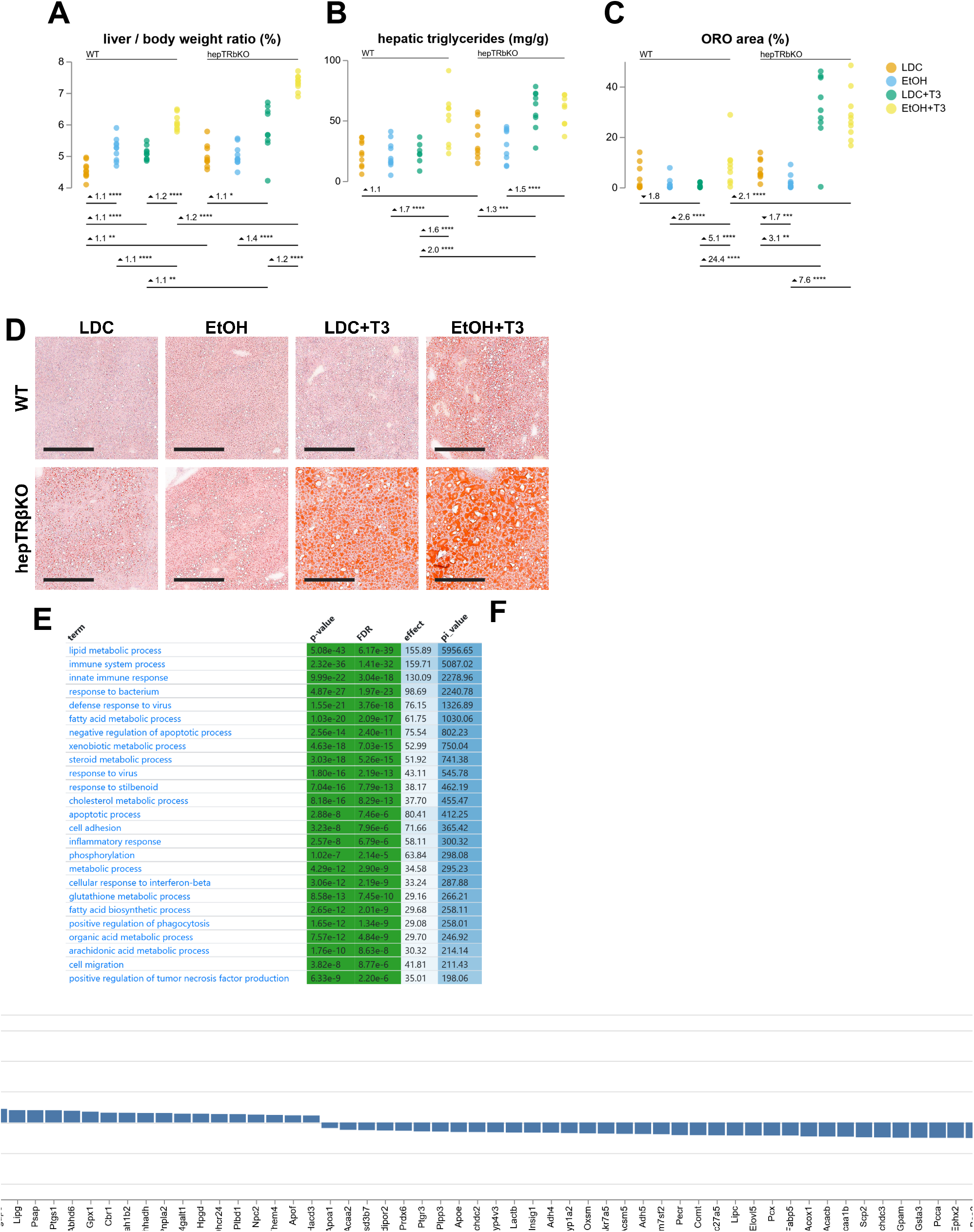
Hepatic fat content of WT and hepTRβKO mice after EtOH and T3 treatment. (A) liver/body weight ratio, (B) hepatic triglycerides and (C) quantification of (D) Oil Red O staining of liver sections (Scale: 300 µm). CFC, BM: *P<0.05,**P<0.01, ***P<0.001, ****P<0.0001. (E) Regulated gene ontology terms (Top 25) from livers of LDC+T3 vs. LDC treated mice, (F) Up- and downregulated genes in gene ontolgy term ‘lipid metabolic process’ from livers of LDC+T3 vs. LDC treated mice. n=7-10.

### Local T3/TRβ signaling stimulates lipid-metabolizing pathways

To determine which genes are influenced by T3 treatment in liver, we performed an RNA-Sequencing analysis, which showed that T3 treatment of WT mice influenced hepatic gene expression in lipid metabolism pathways, including gene ontology terms ‘lipid metabolic process’, ‘fatty acid metabolic process’ and ‘cholesterol metabolic process’ (Figure 2E). Specifically, T3-induced genes included lipoprotein lipase (*Lpl*), which hydrolyzes triglycerides in lipoproteins, the regulatory subunit of the AMP-activated protein kinase (*Prkab1*), which inactivates acetyl-CoA carboxylase (*Acc*) and beta-hydroxy beta-methylglutaryl-CoA reductase (*Hmgcr*), and mitochondrial uncoupling protein 2 (*Ucp2*) (Figure 2F). Repressed were lipogenic genes, such as acetyl-CoA carboxylase (*Acaca*), involved in fatty acid biosynthesis, stearoyl-CoA desaturase (*Scd1*), fatty acid synthase (*Fasn*), ATP citrate synthase (*Acly*) and sterol regulatory element-binding transcription factor 1 (*Srebf1*) (Figure 2F). None of these pathways was found to be enriched after T3 treatment in hepTRβKO mice, neither on control diet nor on EtOH. Thus, gene expression from WT and hepTRβKO mice support the beneficial effect of local T3/TRβ action on hepatic metabolism that clearly contradicts the phenotype of increased hepatic steatosis after T3 treatment. These data suggested that the origin of the detrimental TH effects counteracting the beneficial local TH/TRβ signaling in liver is extrahepatic.

### Hepatic TRβ signaling influences the phenotype of white adipose tissue

Given the seemingly paradoxic results that local hepatic T3/TRβ signaling is metabolically beneficial, but T3 treatment severely aggravated liver steatosis and triglyceride content, we hypothesized that extrahepatic TH effects may be responsible for the latter effect. We therefore studied white adipose tissue (WAT) as a possible source of lipids that accumulate in the liver. Strikingly, WT mice treated with either EtOH or T3 lost more than 50 % of their subcutaneous and epidydimal WAT (scWAT and eWAT) compared to the LDC control mice (Figure 3A). In contrast, hepTRβKO mice lost less WAT by the different treatments compared to their WT littermates.

**Figure 3.**
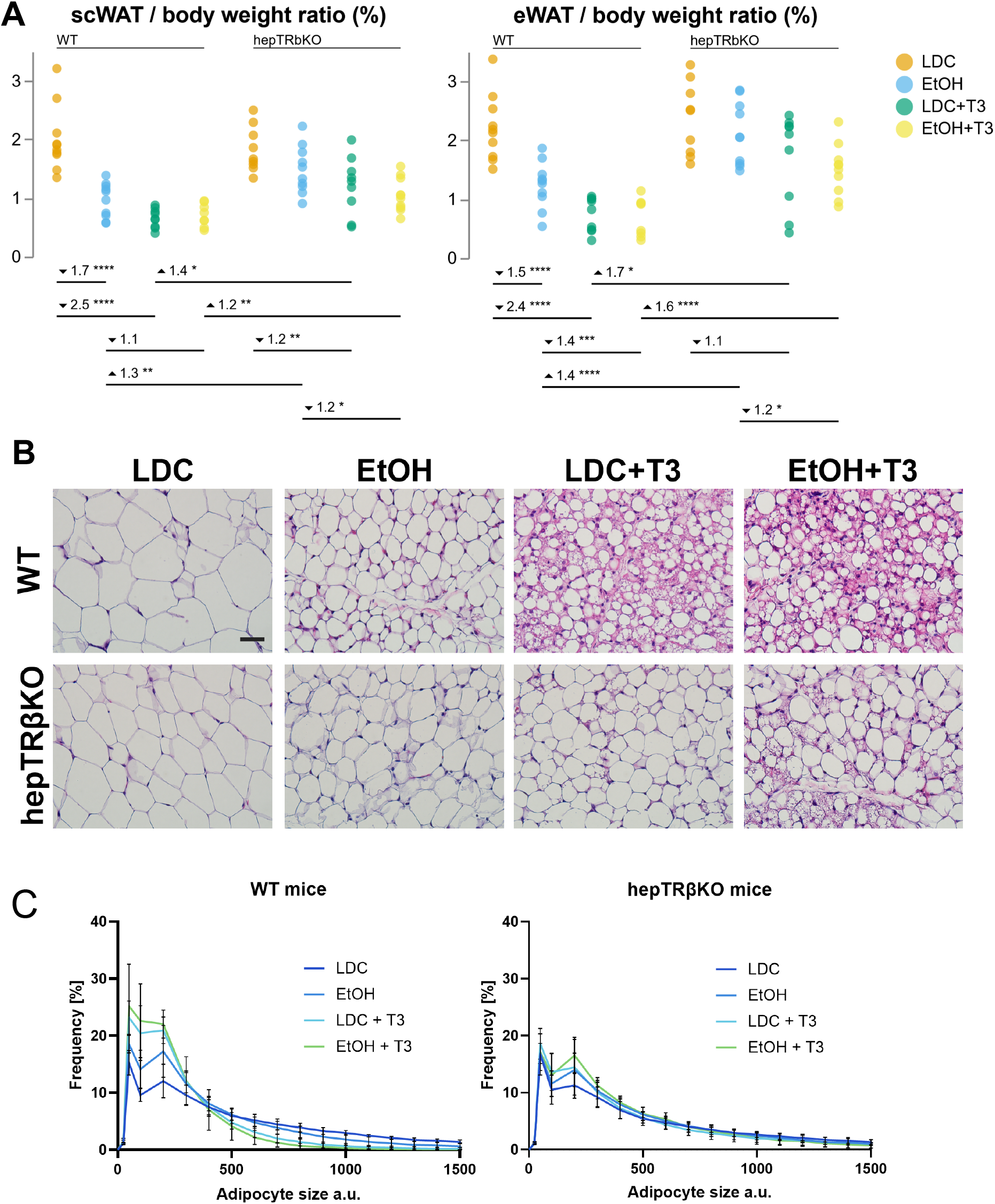
T3-induced adipocyte shrinkage is absent in hepTRβKO mice. (A) scWAT/body weight and eWAT/body weight ratio, (B) hematoxylin & eosin staining of scWAT sections (Scale: 50 µm) and (C) distribution of adipocyte cell size. CFC, BM:*P<0.05, **P<0.01, ***P<0.001, ****P<0.0001. Values are mean±SEM. n=7-10.

We quantified the adipocyte cell size of scWAT (Figure 3B). In WT mice, LDC-treated mice had the largest adipocytes and EtOH led to adipocyte shrinkage. T3 treatment in combination with either diet caused strong adipocyte shrinkage which reached its maximum with EtOH+T3 treatment. Here, the number of smaller adipocytes was increased while the frequency of bigger cells was decreased (Fig. 3C). In hepTRβKO mice, the shrinkage effect by EtOH was maintained. However, the T3-induced effect on adipocytes in WT mice was reduced in hepTRβKO mice independent of the diet and the shift from larger to smaller adipocytes was abolished. These results demonstrate that T3 treatment has strong effects on WAT. Interestingly, given the reduced T3 effect on WAT in mice with TRβ absent only in hepatocytes, hepatic T3/TRβ signaling apparently regulates adipose tissue loss and adipocyte shrinkage.

### Hepatocyte-specific TRβ agonist resmetirom acts in liver but does not affect the pituitary or heart

Resmetirom is a liver-directed TH analog which binds predominantly to TRβ. It could therefore exert beneficial TRβ effects in liver. We compared T3 and resmetirom treatment within the LDC control and EtOH liquid diet. Resmetirom did not change FT3 serum levels (Figure 4A) and effects on FT4 and TT4 levels were only minor compared to T3 treatment. In contrast to T3, resmetirom did not suppress *Tshb* expression in the pituitary (Figure 4B). Systemic T3 treatment was accompanied by cardiac hypertrophy independent of the diet. Again, this effect was absent with resmetirom (Figure 4C). However, serum liver markers AST and ALT were increased by both, resmetirom and T3, independent of the diet (Figure 4D) and blood glucose was reduced by both treatment regimes (Figure 4E). Of note, while TC serum concentrations were decreased by T3 as well as resmetirom, the latter was more potent. Moreover, in EtOH-treated mice T3 treatment resulted in an increase in serum TG concentrations whereas resmetirom treatment reduced serum TG compared to the solvent control. To test whether T3 and resmetirom might act differently on hepatocytes we next performed RNA-sequencing, which showed a high correlation between T3 and resmetirom effects in liver gene expression (Figure 4H). These results confirm that resmetirom acts on the liver but has no effects on the HPT axis and heart. Thus, resmetirom could be more favorable than T3 for treatment in ALD.

**Figure 4.**
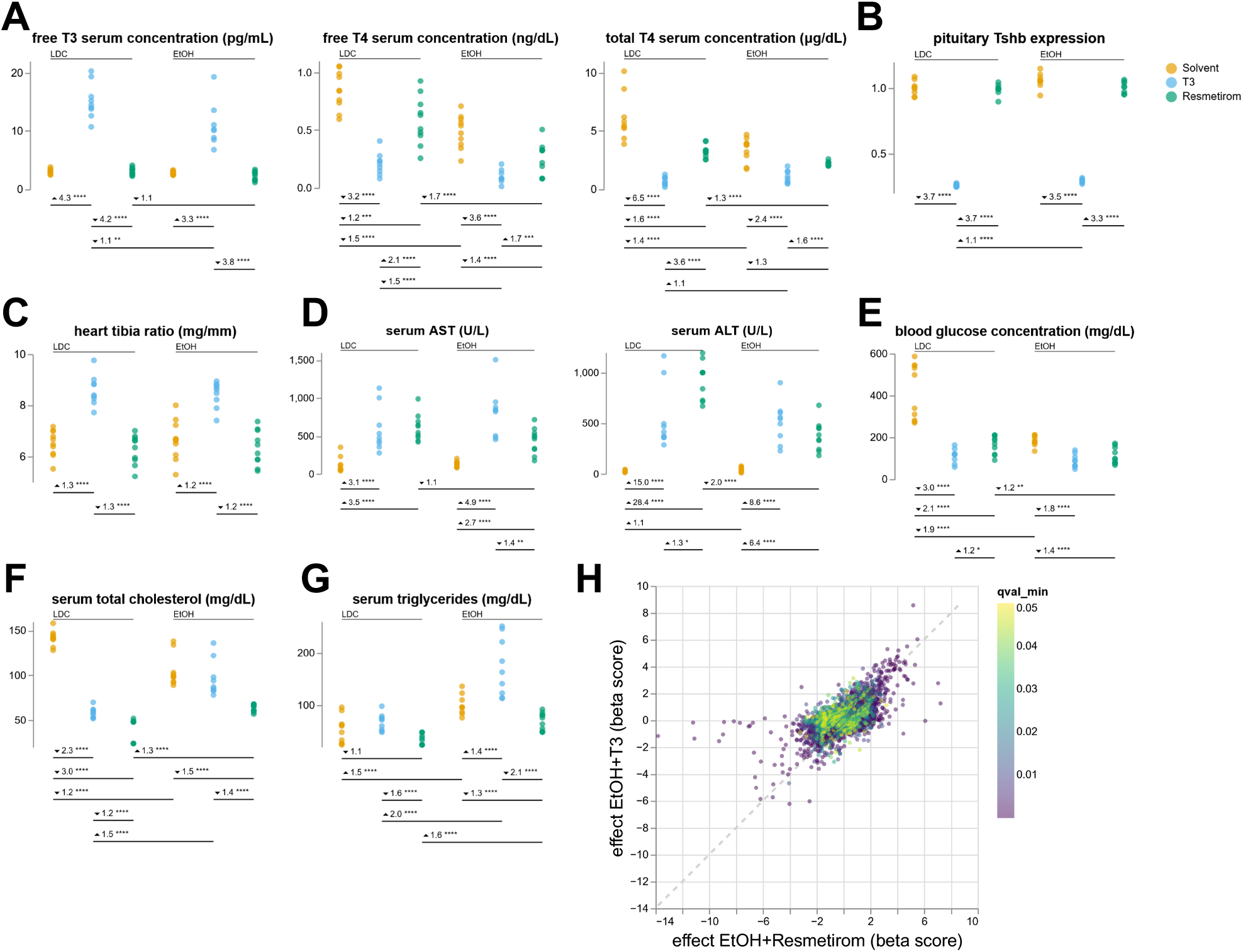
Resmetirom effects on serum parameter in ALD. (A) free T3, free T3 and total T4 serum concentrations, (B) *Tshb* expression in pituitary gland, (C) heart/tibia ratio, (D) serum levels of aspartate transaminase (AST) and alanine transaminase (ALT), (E) blood glucose concentration, (F) total cholesterol serum concentrations and (G) serum triglyceride levels. CFC, BM: *P<0.05, **P<0.01,***P<0.001, ****P<0.0001. (H) DEG comparison of EtOH+T3 vs. EtOH and EtOH+Resmetirom vs. EtOH. n=7-10.

### Resmetirom induces extrahepatic effects in WAT and does not improve ethanol-induced liver steatosis

We compared resmetirom and T3 treatment in the ALD model. We observed for both diets, LDC and EtOH, that in resmetirom-treated mice the liver/body weight ratios were not different from solvent controls but lower than those of T3-treated mice (Figure 5A). Unlike for T3, hepatic TG content was reduced with resmetirom and not increased by resmetirom on EtOH diet compared to solvent control. But TG content with EtOH + resmetirom was increased compared to resmetirom alone (Figure 5B). This was also seen in Oil Red O staining of liver sections (Figure 5 C, D).Next, we studied whether resmetirom has any effect on WAT. Already on LDC diet, resmetirom reduced the scWAT/body weight and eWAT/body weight ratios to half, very similarly to T3 (Figure 5F). EtOH diet did not further increase WAT loss in the treatment groups. Adipocyte histology showed that resmetirom shifted the cell size of adipocytes towards smaller cells sizes, although less than T3 (Figure 5G, H).For a liver-directed TRβ agonist such extrahepatic effects of resmetirom were quite unexpected. Therefore, we studied whether resmetirom induced gene expression changes in scWAT similar to T3, by comparing RNA-sequencing data from liver and WAT for T3 and resmetirom on EtOH diet in WT mice. T3 and resmetirom had similar effects on gene expression in liver (Fig. 5E). But in contrast to T3, resmetirom did not influence WAT gene expression (Figure 5I). This suggests that resmetirom is indeed a liver-directed thyromimetic and that, consequently, the observed WAT loss induction by resmetirom is not a direct effect in WAT but may originate in liver.

**Figure 5.**
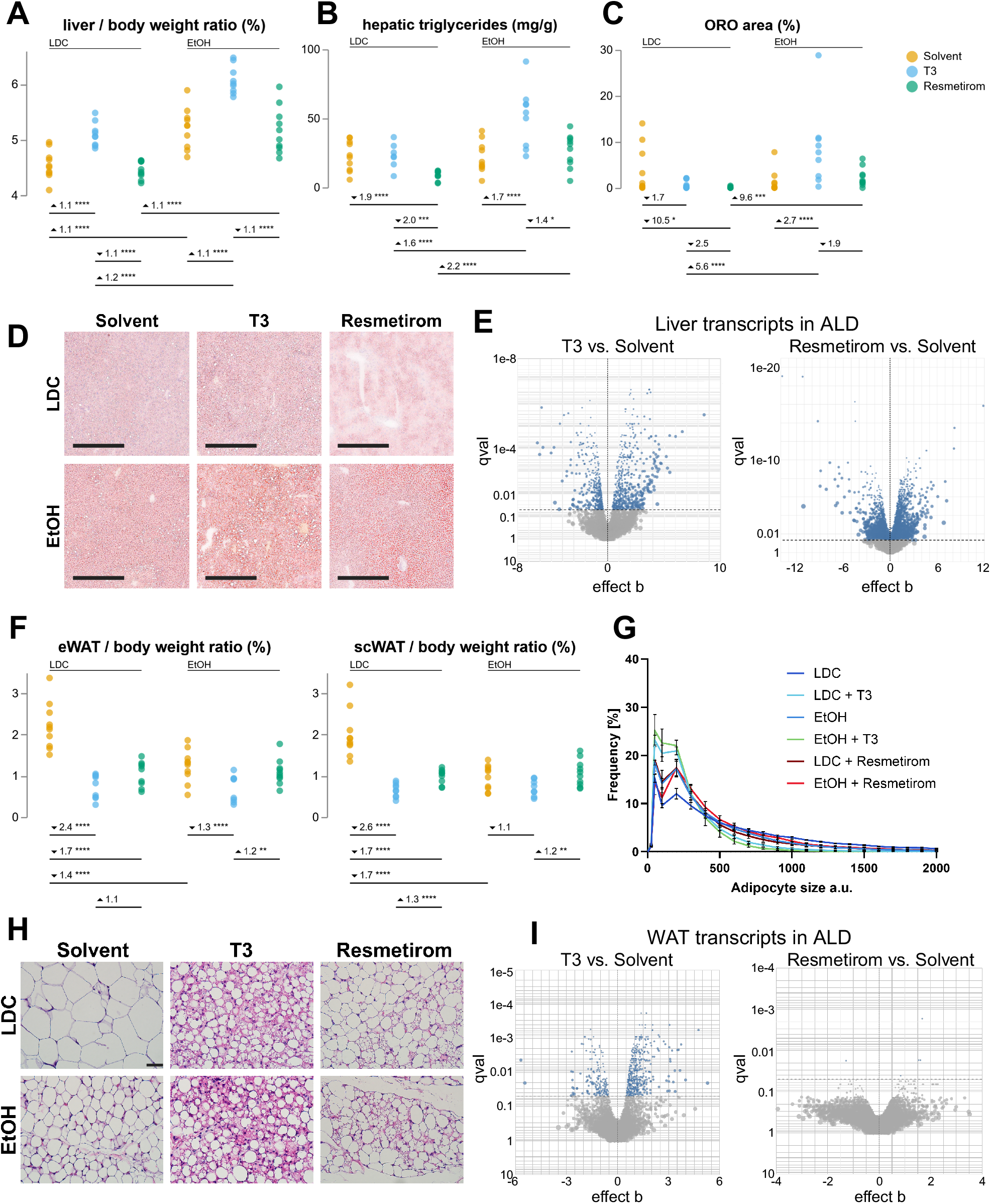
Resmetirom effects on hepatic steatosis and WAT in ALD. (A) liver/body weight ratio, (B) hepatic triglycerides, (C) quantification of (D) Oil Red O staining of liver sections (Scale: 300 µm), (E) liver transcripts of T3 vs. solvent and resmetirom vs. solvent in ALD, (F) scWAT/body weight and eWAT/body weight ratio,(G) distribution of adipocyte cell size and (H) hematoxylin & eosin staining of scWATsections (Scale: 50 µm) and (I) WAT transcripts of T3 vs. solvent and resmetirom vs. solvent in ALD. CFC, BM: *P<0.05, **P<0.01, ***P<0.001, ****P<0.0001. Values are mean±SEM. n=7-10.

## Discussion

ALD is characterized by fat accumulation in hepatocytes and multiple mechanisms contribute to this hepatic steatosis. Major mechanisms are the disruption of mitochondrial fatty acid β-oxidation and migration of lipids to the liver from extrahepatic organs.^22^ THs promote hepatic catabolism of lipids and reduces liver lipid content.^23^ Given these beneficial effects of TH and the liver-directed TRβ analog resmetirom, we hypothesized that TH and resmetirom could also be beneficial in ALD, in which steatohepatitis is a key driver of mortality. Surprisingly, TH treatment severely aggravated EtOH-induced hepatic steatosis in WT mice, although the hepatic TRβ still exerted the known beneficial effects on hepatic lipid metabolism. These apparently contradicting results could be explained by the finding of extrahepatic action of TH and resmetirom: both induced severe WAT loss, presumably leading to flooding of the liver with lipids, which could not be compensated due to the EtOH diet. Mathur *et al*. recently showed in mice that EtOH consumption stimulates adipose lipolysis.^24^ Thus, WAT lipolysis critically contributes to hepatic steatosis and oxidative stress during ALD development. Yet, neither T3 nor EtOH treatment alone increased hepatic steatosis, but both treatments together resulted in severe steatosis. Apparently, lipids mobilized by T3 or EtOH could be metabolized in the liver. But their combination, adding the disruption of hepatic fatty acid β-oxidation by EtOH, exceeded the compensatory ability of the liver. EtOH and TH act synergistically on WAT loss and lipid flooding of the liver, resulting in the observed exacerbation of hepatic steatosis on EtOH diet and TH.

While WAT reduction with release of lipids and their accumulation in the liver could explain aggravation of hepatic steatosis by T3 in ALD, the mechanism is less obvious. A surprising finding was that absence of TRβ only in hepatocytes (hepTRβKO mice) diminished T3 effects in WAT compared to WT mice: The T3-induced WAT loss seen with EtOH or T3 was reduced in hepTRβKO mice and T3-induced WAT adipocyte shrinkage was almost completely abrogated in hepTRβKO mice. Thus, hepatic TRβ action was necessary to induce WAT loss.

Resmetirom did not influence WAT gene expression. Still, resmetirom induced WAT loss of about 50%, comparable to T3. Resmetirom is considered a hepatocyte-specific TRβ agonist but exerted strong extrahepatic effects on scWAT. The important question is now: How can presence or absence of TRβ in hepatocytes influence the phenotype of WAT? Recently, the concept has been established thatthe liver acts like an endocrine organ and regulates metabolism via secreted proteins, called hepatokines, that signal to distant tissues, e.g. to WAT.^25^ Examples are FGF21 and ANGPTL4. We hypothesize that in hepatocytes T3 and resmetirom induce a hepatokine via TRβ that communicates to WAT, constituting a novel T3/TRβ-liver-WAT axis. To fully understand the extrahepatic role of hepatic TH/TRβ action in general physiology and in ALD, the hepatokine needs to be identified and the T3/TRβ-liver-WAT axis studied in detail. TH effects in one organ (WAT) that originate from T3/TRβ action in another organ (liver) also expand the concept of local TH action and warrant further clarification.

What is the clinical relevance of these results? Resmetirom is a hepatocyte-specific TRβ analog and has been developed with the idea to harness the beneficial TH effects in hepatocytes and avoid extrahepatic adverse effects. We found earlier that resmetirom is TRβ-selective and requires the hepatocyte-exclusive transporter OATP1B1 for transport into cells.^7^ A phase III study (MAESTRO-NASH) has been completed in December 2022 and the FDA criteria for NASH resolution have been met.^8^ Because resmetirom demonstrated substantial improvement over available therapy on clinically relevant endpoints, resmetirom was conditionally approved by the FDA and the EMA for treatment of MASH. Yet, the phase III study data showed that histological NASH resolution was observed only in 26% (80 mg) and 30% (100 mg) with resmetirom vs. 10% with placebo. Response rates were even lower for fibrosis improvement (24% and 26% vs. 14%, respectively). Thus, despite being considered a breakthrough therapy, most patients (≥70%) did not fully respond to resmetirom. The clinically relevant question is why more than two thirds of patients did not respond. Based on our results, resmetirom as a hepatocyte-specific TRβ analog induces WAT loss and probably release of fatty acids that are received by the liver, which could diminish resmetirom’s efficacy. We hypothesize that resmetirom, like T3, induces a hepatokine, stimulating WAT lipolysis and increasing hepatic TG, especially when the lipolytic effect in WAT is augmented by EtOH. This liver-WAT axis and its aggravation by EtOH may explain the low response rate of resmetirom in many NASH patients, because alcohol consumption is currently not completely prohibited in NASH and alcohol consumption and high calory intake are frequent co-risk factors. Harmful alcohol consumption in presumed NASH has been reported.^26^ This hypothesis has clinical and therapeutical relevance for application of resmetiromand alcohol consumption in patients with MASH. Resmetirom may be much more efficient in resolving liver steatosis in ALD with cessation of alcohol consumption.In summary, we show *in vivo* that TH and EtOH synergistically increase WAT loss and hepatic steatosis. This effect is in part remotely regulated by hepatic TRβ, constituting a novel TH-liver-WAT axis, and may diminish resmetirom’s therapeutic efficacy.

## Acknowledgments

We are grateful for the continued dedicated support from Prof. Dr. G. Hilken, Dr. A. Wißmann, Dr. P. Dammann, Dr. M. Dubicanac and the staff of the animal facility at the University Hospital Essen. We thank Andrea Jaeger, Dorothe Möllmann and Martin Schlattjan for their dedicated technical assistance. The graphical abstract was created with BioRender.com.

## Figure Legends

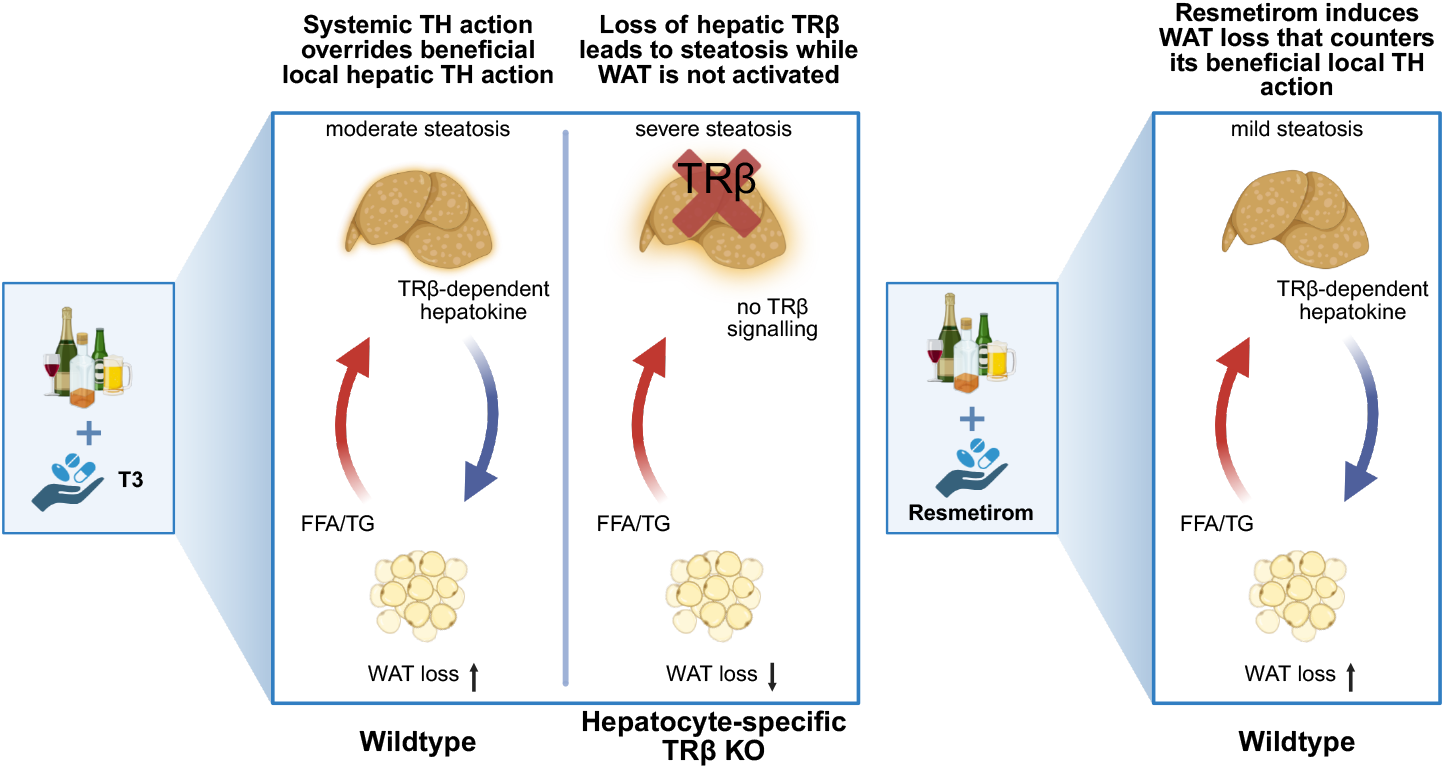

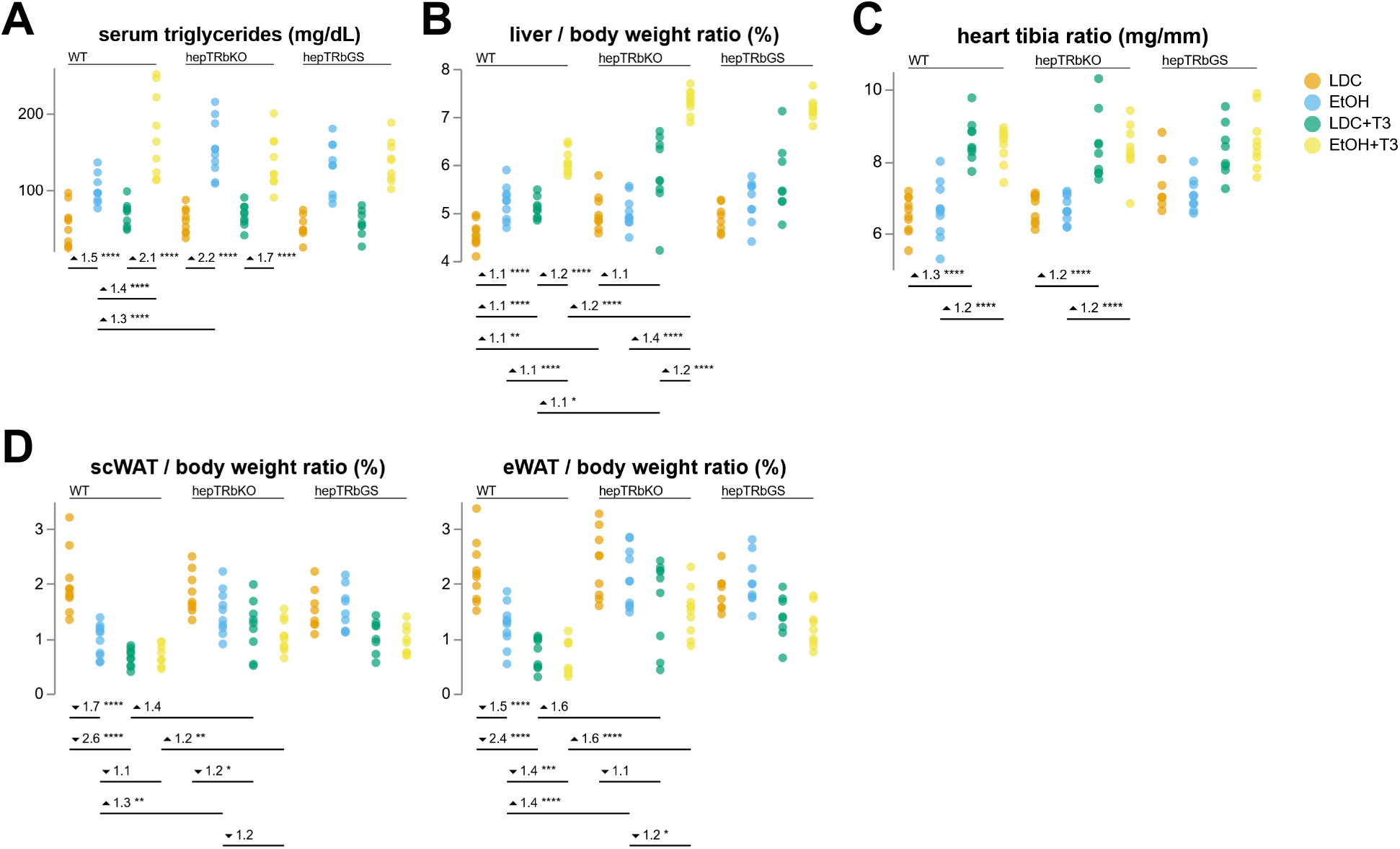

